# Multi-layered network-based pathway activity inference using directed random walks: application to predicting clinical outcomes in urologic cancer

**DOI:** 10.1101/2020.07.22.163949

**Authors:** So Yeon Kim, Eun Kyung Choe, Manu Shivakumar, Dokyoon Kim, Kyung-Ah Sohn

## Abstract

**Motivation:** To better understand the molecular features of cancers, a comprehensive analysis using multi-omics data has been conducted. Additionally, a pathway activity inference method has been developed to facilitate the integrative effects of multiple genes. In this respect, we have recently proposed a novel integrative pathway activity inference approach, iDRW, and demonstrated the effectiveness of the method with respect to dichotomizing two survival groups. However, there were several limitations, such as a lack of generality. In this study, we designed a directed gene-gene graph using pathway information by assigning interactions between genes in multiple layers of networks.

**Results:** As a proof-of-concept study, it was evaluated using three genomic profiles of urologic cancer patients. The proposed integrative approach achieved improved outcome prediction performances compared with a single genomic profile alone and other existing pathway activity inference methods. The integrative approach also identified common/cancer-specific candidate driver pathways as predictive prognostic features in urologic cancers. Furthermore, it provides better biological insights into the prioritized pathways and genes in an integrated view using a multi-layered gene-gene network. Our framework is not specifically designed for urologic cancers and can be generally applicable for various datasets.

**Availability:** iDRW is implemented as the R software package. The source codes are available at https://github.com/sykim122/iDRW.

## 1 Introduction

To better understand the complex biological mechanism underlying cancer progression and prognosis, a comprehensive analysis using multi-omics data has attracted great attention to reveal the distinctive and shared molecular features of cancers. Many multi-omics studies have been conducted to discover novel biomarkers associated with cancers and predict clinical outcomes precisely (Huang *et al*., 2017; Kim *et al*., 2014; Lee *et al*., 2017; Shivakumar *et al*., 2017; Sohn *et al*., 2013; El-Manzalawy *et al*., 2018; Kim *et al*., 2015b, 2012). For a comprehensive analysis of multi-omics data, it is crucial to understand the complex interplay between genes across different omics layers. To utilize the interaction effect between genes across multi-omics data, network-based integrative approaches have several advantages, such as utilizing the interrelationships among multi-omics data, better biological interpretation, and improved outcome prediction power, as shown in many studies (Jeong *et al*., 2015; Kim *et al*., 2015a; Lee *et al*., 2019; Vangimalla *et al*., 2016; Wang *et al*., 2017; Kim *et al*., 2017).

To effectively combine different types of genomic features on the graph, most network-based integrative methods have focused on incorporating prior knowledge such as pathway or subtype information in many cancer studies (Hung and Chiu, 2017; Liu *et al*., 2015; Dimitrakopoulos *et al*., 2018). Biological pathways contain interactions among molecules in a cell, and enormous amounts of information on pathways and interactions are readily available in many pathway databases. To identify biologically meaningful molecular features and investigate interaction effects among them, many pathway-based approaches have been proposed based on the network structure (Hu *et al*., 2017; Stoney *et al*., 2018). In this respect, pathway activity inference methods have been developed to produce pathway-level features and corresponding activity scores for robust and accurate prediction and better interpretation. The pathway activity score can be simply computed with summary measures of gene sets, which take the arithmetic mean or the median of the gene expression values of the pathway member genes (Guo *et al*., 2005). Lee, et al. proposed a precise disease classification model by inferring pathway activities for each patient (Lee *et al*., 2008). The pathway activity is defined as the summarized gene expression levels of its condition-responsive genes (CORG), which are the subset of genes in the pathway for which the combined expression shows optimal discriminative power for the disease phenotype. Tomfohr, et al. proposed a pathway-level analysis of gene expression (PLAGE), which derived activity scores from a vector of the singular value decomposition of the given gene set (Tomfohr *et al*., 2005). Additionally, many other pathway activity inference approaches have been proposed in different cancers or other complex diseases (Temate-Tiagueu *et al*., 2016; Wang *et al*., 2019). Those pathway activity inference methods simply take pathways as the set of genes and summarize the gene expression levels; thus, the interaction effects between genes are not considered. In this respect, several studies utilized gene interactions based on network structure. A denoising algorithm based on relevance network topology (DART) derived perturbation signatures that reflect gene contributions in each pathway on the relevance network for improved pathway activity inference (Jiao *et al*., 2011). Liu, et al. proposed a directed random walk-based pathway activity inference method (DRW) to consider the topological importance of the genes on the network that can be highly associated with diseases (Liu *et al*., 2013). DRW has been extensively studied with many variations, including DRW based on a gene-metabolite graph and DRW for survival prediction (Liu *et al*., 2017, 2015).

However, most existing pathway activity inference methods targeted a single genomic profile alone. In this respect, we have recently investigated the effectiveness of the network-based integrative pathway activity inference method for multi-omics data integration (iDRW) (Kim *et al*., 2019, 2018). One of the limitations of previous studies on iDRW lies in the lack of a comprehensive analysis of different levels of genomic data. The integrated gene-gene network in iDRW was formally designed specifically based on the data structure, resulting in a lack of generality. Due to the complexity of the multi-omics network, multiple network scenarios should be considered. Furthermore, it was validated in a classification model that divides long-term and short-term survival groups, not survival prediction. As there are no clear criteria for dichotomizing two survival groups, it highly depends on the data.

To overcome those limitations, we propose a general framework for integrative pathway activity inference on the multi-omics network and investigate multiple network scenarios. To reflect the interaction effects of genes, we designed a directed gene-gene graph in multiple layers by assigning within-layer interactions and between-layer interactions considering multiple scenarios. We inferred pathway activities by performing a random walk with restart (RWR) on the multi-layered network. As a result, iDRW transforms the multiple genomic profiles into a single pathway profile on the graph. The inferred pathway profile is validated with the outcome prediction models. We prioritized pathways, visualized the multi-omics network and extensively analyzed the pathway activity patterns.

As a proof-of-concept study, the proposed method is applied for the integrative analysis of urologic cancer. Urologic cancers include prostate, kidney and bladder cancer that share a common genetic architecture across different types. Here, we considered two types of outcome prediction models (overall survival days and regional lymph node or distant metastasis) for bladder and kidney cancer. The proposed method selects cooperative potential driver pathways associated with clinical outcomes. We also provide extensive analyses of distinguishable and shared molecular features across two different cancers. The overview of the integrative urologic analysis using the proposed method is illustrated in **Figure 1**. The main contributions of this study are summarized as follows.

- We propose a general framework for integrative pathway activity inference on the multi-omics network and investigate multiple scenarios of the multi-layered gene-gene graph.
- We validated our framework with the integrative network-based analysis of urologic cancers using three types of genomic profiles considering two types of clinical outcomes.
- We performed a sophisticated pathway-based integrative analysis: inferred pathway activity pattern analysis, pathways prioritization, and multi-omics network visualization.

**Fig. 1.**
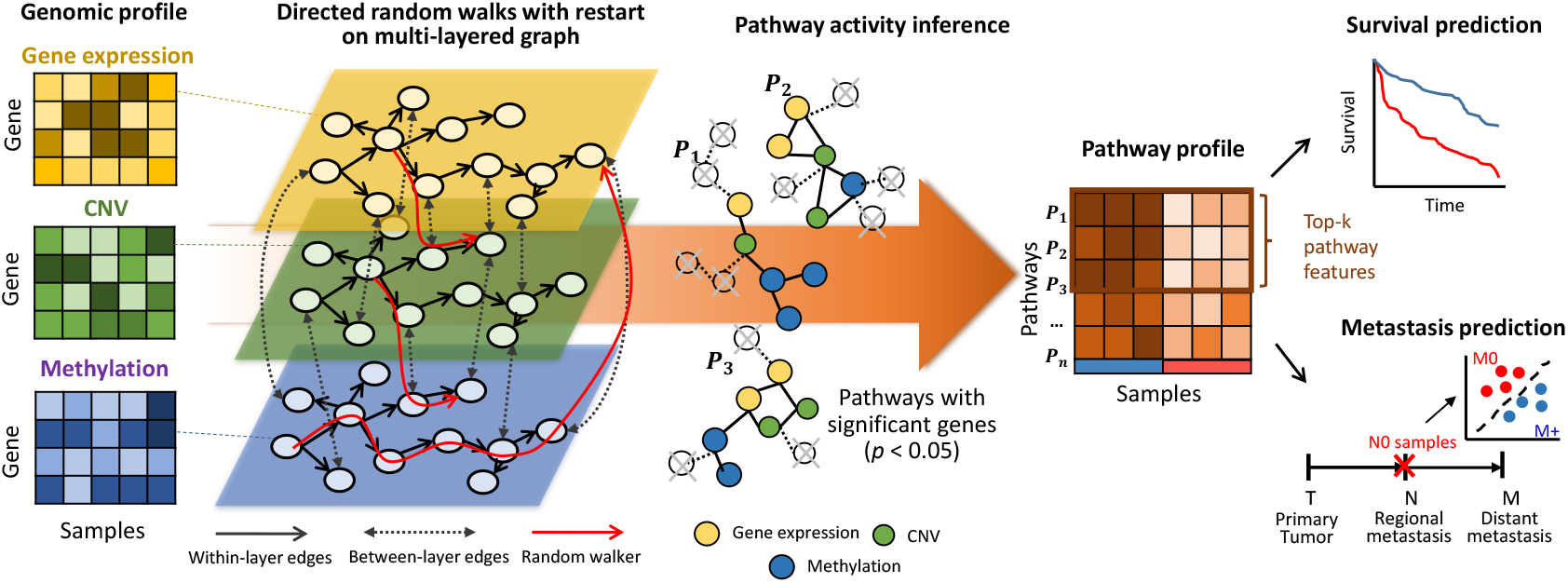
Overview ofan integrative directed random walk-based pathway activity inference on the multi-layered gene-gene graph (iDRW). The iDRW was applied for the integrative urologic cancer analysis using three genomic profiles. The genes of each genomic profile are represented as nodes on each layer of the graph. A directed random walk with restart (RWR) is performed on the multi-layered graph and the gene weights are iteratively updated based on the graph structure. Pathway activities are inferred using the subset of pathway member genes that are significantly associated with the outcome (*p* < 0.05) by combining the normalized gene expression values, a statistical score of genes that represents the statistical significance, and the updated gene weights by performing RWR on the multi-layered graph. For a systematic view, iDRW transforms the multiple genomic profiles into a single pathway profile. The pathway profile is used as an input to train the prediction model. The framework was evaluated for survival and metastasis prediction performance and prioritization of the top-k pathways for cancer prognosis and metastatic progression.

## 2 Methods

### 2.1 Pathway-based multi-layered gene-gene graph

Let the multi-layered gene-gene graph **G** = (V, E, X) be composed of *L* layers. A gene-gene graph on the *i*-th layer is defined as **G**_*i*_, = (V_*i*_, E_*i*_, X_*i*_), where V_*i*_ is a set of nodes (genes), E_*i*_ is a set of directional edges, and 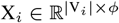 is a feature matrix of the nodes at the *i*-th layer. Then, 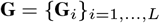.

Let *v* ∈ V_*i*_, *w* ∈ V_*j*_ be a node of the graph on the *i, j*-th layer, *e_vw_* = (*v, w*) ∈ E be an edge between the node *v* and *w*, and 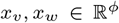 feature vectors of the nodes *v* and *w*. In this study, X_*i*_ is a genomic profile of a gene-by-sample matrix and *x* is the *ϕ*-sized gene expression vector. E is composed of within- and between-layer interactions. The within-layer interactions are derived from the pathway-based gene-gene interactions. The set of pathways P are obtained from pathway databases such as KEGG, Reactome, and WikiPathways, i.e. P = P_1_ ∪ ⋯ ∪ P_*N*_ for *N* pathways. Then, P = (M, I) where M are molecules (genes) and I are molecular interactions of genes, defined in the pathway database. For each layer, the nodes and edges of **G**_*i*_ were derived from P:

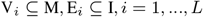

We define between-layer interactions with two possibilities. We assign bi-directional edges ^1)^ between all pairwise combinations of *v* and *w*:

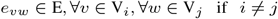

or ^2)^ if there is a higher correlation between nodes *v* and *w* than a threshold, i.e., if |*corr*(*x_v_, x_w_*)| ≥ *θ*, where |*corr*(*x_v_, x_w_*)| represents the magnitude of the Pearson correlation coefficient between feature vectors of node *v* and *w* (correlation of the gene expressions between gene *v* and *w*) in two different layers, and *θ* is a threshold value of the correlation. We set the threshold value *θ* to 0.5, as it is generally accepted to have a low correlation when it is lower than 0.5 (Hinkle *et al*., 2003). The adjacency matrix *A* of the multi-layered graph **G** is derived as *A_vw_* = 1 if *e_vw_* ∈ E and 0 otherwise. We note that *A* is an asymmetric matrix because **G** is a directed graph, i.e., *A* ≠ *A*^T^.

### 2.2. Directed random walks with restart on the multi-layered graph

To infer pathway activities from multiple genomic profiles, we performed random walk with restart (RWR) on the multi-layered directed gene-gene graph **G**. The purpose of performing RWR is to update genes considering their interaction effects within- and between-genomic layers and transform them into a pathway-level activity matrix. It should be noted that this step was performed using the entire dataset, and the transformed pathway profile (pathway-by-sample matrix) was used as an input to the training model. In the training phase, the samples were divided into training and validation sets for cross-validation. A set of genes, so-called seeds, the starting points of the random walk algorithm, was used to explore the neighborhood and iteratively update the nodes of the graph. To start a random walk on a graph, we initialized the seed genes by univariate statistical analysis to assess the significant association between each gene and the clinical outcome. For each node (gene) *v*, we obtained a statistic score (*z_v_*) and *p*-value of the statistical significance of the model (*p_v_*) using a feature vector *x_v_*. The method took three kinds of the outcome variables: survival time, binary and multi-class outcomes. To measure the significant associations of genes with survival time, univariate cox regression analysis was performed for each gene, stratified by several confounding factors including age, gender, and TNM stage, using a cox proportional hazards regression model (Andersen and Gill, 1982). We assessed the Wald statistic value as a statistic score and *p*-value corresponding to the ratio of each gene’s regression coefficient to its standard error. For binary or multiclass outcome, it performed a two-tailed t-test or an analysis of variance (ANOVA) for each gene to test the significant differences between the group means of each class. This process produced a *t* or *F* value as a statistic score and a *p*-value for each gene. The initial weight vector 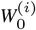 for the *i*-th layer is formally defined as:

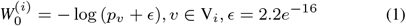

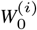 is normalized to scale the range between 0 and 1 and combined to develop 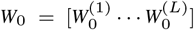. Then, *W*_0_ is *l*_1_-normalized to a unit vector. A random walker starts on a source node *s* (seeds) and transits to a randomly selected neighbor or returns to the source node *s* with a restart probability *r* at each time step *t*. *W*^(*t*)^ is iteratively updated with:

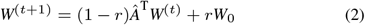

where *A*^(*t*)^ is the weight vector in which the *i*-th element represents the probability of being at node *i* at time step *t, r* is the restart probability, and *Â* is a row-normalized matrix of the adjacency matrix *A* of the multilayered gene-gene graph **G**. We set the restart probability *r* to 0.3 as it has been shown that the performance is not sensitive to the varying *r* (Liu *et al*., 2013). After a number of iterations, it is guaranteed to converge to a steady state W when |*A*^(*t*+1)^ – *A*^(*t*)^| < 10^−10^, as previously shown (Liu *et al*., 2017, 2013, 2015). The final weight vector of nodes in the multi-layered graph **G** was obtained as 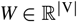.

### 2.3 Pathway activity inference

We inferred pathway activities with the set of statistically significant genes. Let the *i*-th pathway P_*i*_ include *n_i_* number of genes that are significantly associated with the outcome (*p*-value < 0.05), and *v_k_* be the *k*-th significant gene, i.e., *v_k_* ∈ P_*i*_, *k* = 1,⋯, *n_i_*. The pathway activity *PA_i_* for the *i*-th pathway is defined as:

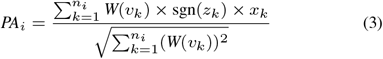

where *W*(*v_k_*) is the final weight of node *v_k_* (gene) which was updated from **Equation** (2); *x_k_* is the node feature vector (vector of gene expression values from the original genomic data); *z_k_* is the statistic score derived from the univariate statistical analysis (cox regression/t-test/ANOVA); and sgn(*z_k_*) is the sign of the statistic score indicating a positive or negative correlation between the gene expression values and clinical outcome. This formula makes the pathway activity score low when it is combined with negatively correlated genes with the risk of patients. For each pathway, the pathway activity is computed across all samples, considered as a pathway profile, i.e., 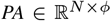. The high value of the pathway activity score indicates that the corresponding pathway highly affects the risk of patients. Finally, iDRW combines the feature matrices in *L* layers into a single pathway profile based on the multi-layered graph as 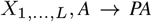 where 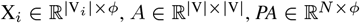.

## 3 Experiments

### 3.1 TCGA urologic cancer datasets

We obtained RNA-Seq gene expression, copy number variation, and DNA methylation profiles of the TCGA bladder cancer (BLCA) and kidney clear cell carcinoma (KIRC) dataset. Gene expression data were measured using Illumina HiSeq 2000 RNA Sequencing, which is level 3 data from the TCGA data coordination center. It consisted of 20,530 genes, which are gene-level transcription estimates, as in the log-transformed RSEM normalized count. The gene-level copy number variation (CNV) data were estimated using the GISTIC2 method, which consisted of 24,776 genes. RNA-Seq geneexpressionandCNV datawere downloaded from the UCSCXena platform (Goldman *et al*., 2019). DNA methylation data were obtained as a gene-level feature by selecting the probe having a minimum correlation with the expression data for each gene from the Broad Institute GDAC Firehose (GDAC, 2016). In this study, the overlapping 16,904 genes and 400 patients were in BLCA data across three genomic profiles. Likewise, 17,125 genes and 313 patients were in KIRC data.

Although there are 6 types of urologic cancer datasets in TCGA, we excluded the testicular cancer (TGCT), kidney chromophobe (KICH), dataset due to the extremely small number of cases, which were 134 and 65, respectively. The kidney papillary cell carcinoma (KIRP) and prostate cancer (PRAD) dataset had a sufficient number of samples; however, those cancers were excluded due to the excessive rate of censoring, which was 85.1% and 98.2%. Excessive censoring rate leads to the risk of bias to the prediction model and severely harms the model performance and interpretation of the results (Zhu *et al*., 2017).

### 3.2 Data preprocessing

There were 811 and 756 missing values in the DNA methylation data for BLCA and KIRC, respectively. We imputed them with a median of the corresponding patient’s data. We excluded patients whose clinical outcome variables were not recorded or inaccurate such as the negative values of survival days.

For each cancer dataset, overall survival (OS), event status, age, gender, and TNM stage were used as clinical variables. The event status is a binary variable with the event occurred (1) and right-censored (0). There were 223 censored and 173 uncensored samples in BLCA (censoring rate: 56.3%), and 209 censored and 102 uncensored samples in KIRC (censoring rate: 67.2%). The clinical variables were dichotomized as ages into 0 (< 65 years) or 1 (≥ 65 years), T stages into 0 (T0-2) or 1 (T3-4), N stages into N0 or N1-3 (N+), and M stages into M0 or M1. We filled some of the unknown pathologic stages based on the American joint committee on cancer (AJCC) staging system. In addition, the missing N or M stages were filled according to the number of lymph nodes that were positive or metastatic sites. For example, if the number of lymph nodes positive was greater than 0, they were categorized as N+. If the metastatic site was recorded as ‘lymph node only’, they were regarded as M0. Metastatic site features were recorded only in the BLCA dataset.

We considered OS and three types of metastasis prediction models; the model to predict the risk of patients with any metastasis (any T/ N+/M1); patients with regional lymph node metastasis without distant metastasis (any T/N+/M0); patients with distant metastasis without regional lymph node metastasis (any T/N0/M1). Due to the number of samples for each class, we evaluated our model with any or regional metastasis in BLCA, and any or distant metastasis prediction model in the KIRC dataset. The total number of samples for each clinical feature are shown in **Table 1**.

**Table 1.** Summary statistics of clinical features in the TCGA bladder cancer (BLCA) and kidney clear cell carcinoma (KIRC) data

| Data type |  | BLCA | KIRC |
| --- | --- | --- | --- |
| Number of samples |  | 400 | 313 |
| Age | < 65 years | 147 (36.8%) | 193 (61.7%) |
| | ≥ 65 years | 253 (63.2%) | 120 (38.3%) |
| Gender | Male | 295 (73.8%) | 201 (64.2%) |
|  | Female | 105 (26.2%) | 112 (35.8%) |
| Stage T | T0-2 | 148 (37.6%) | 196 (62.6%) |
|  | T3-4 | 246 (62.4%) | 117 (37.4%) |
| Stage N | N0 | 261 (67.4%) | 244 (87.8%) |
|  | N+ | 126 (32.6%) | 34 (12.2%) |
| Stage M | M0 | 340 (86.1%) | 258 (82.7%) |
|  | M1 | 55 (13.9%) | 54 (17.3%) |
| Overall survival (OS) | Survival days | 810.5±833.8 | 1310.3±1062.7 |
|  | Uncensored | 173 (43.7%) | 102 (32.8%) |
|  | Censored | 223 (56.3%) | 209 (67.2%) |
| Any metastasis | Positive (any T/N+/M1) | 51 | 26 |
|  | Negative (any T/N0/M0) | 260 | 216 |
| Regional metastasis | Positive (any T/N+/M0) | 70 | - |
|  | Negative (any T/N0/M0) | 260 | - |
| Distant metastasis | Positive (any T/N0/M1) | - | 28 |
|  | Negative (any T/N0/M0) | - | 216 |

### 3.3 Multi-layered network construction

We investigated the integrative effect of the iDRW method with a comparison of the single-layered graph, which corresponds to the DRW method, for each genomic profile, denoted as **G**^E^, **G**^C^, and **G**^M^. To investigate the prognostic effect of combining each genomic profile, we experimented with all possible combinations between each layer, denoted as **G**^EC^, **G**^EM^, **G**^CM^, and **G**^ECM^. For example, **G**^EC^ = {**G**^E^, **G**^C^} is a two-layered gene-gene graph combining the gene expression and CNV profile. We experimented with four scenarios of constructing **G**^ECM^: the one that assigns within-layer edges based on pathway-based gene-gene interactions to^1)^ all genomic profiles or ^2)^the gene expression profile only to demonstrate the effect of combining them with the copy number variation (CNV) or DNA methylation profile. For each of the former scenarios, we assigned between-layer edges from ^a)^all pairwise combinations of genes or ^b)^a pair of genes with a correlation coefficient of the expression value greater than 0.5.

To assign pathway-based gene-gene interactions to within-layer edges for each layer, we constructed a pathway-based directed gene-gene graph using the KEGG pathway database (Kanehisa and Goto, 2000). We parsed KGML (KEGG XML) files of 327 KEGG pathways into graph models using the R package, KEGGgraph (Zhang and Wiemann, 2009). For each pathway, we included all the nodes and edges where the node type is a gene. The genes in the pathway were annotated as the respective HUGO gene symbols. We merged 327 human pathways into a pathway-based genegene graph. In total, 7,390 nodes and 58,426 edges were obtained. For each cancer dataset, we had three types of genomic profiles: RNA-Seq gene expression, CNV, and DNA methylation profile, which were considered as three layers. For each layer, the overlapping genes between genes from the genomic profile and pathway-based gene-gene graph were present: |V^E^| = 6,827 (BLCA & KIRC), |V^C^| = 7,077 (BLCA & KIRC), |V^M^| = 5,805 (BLCA) and 5,894 (KIRC). Each genomic profile was normalized for the mean to be 0 and standard deviation to be 1 across all the samples.

To demonstrate the effectiveness of the DRW-based integrative approach, we additionally compared iDRW with three state-of-the-art pathway activity inference methods for each genomic profile: CORG (Lee *et al*., 2008), PLAGE (Tomfohr *et al*., 2005), and DART (Jiao *et al*., 2011). CORG and PLAGE were implemented with the R package GSVA, with default settings (Hanzelmann *et al*., 2013). As each of those pathway activity inference methods has been previously developed based on a single genomic profile, we assessed pathway activities across samples from **G**^E^, **G**^C^, and **G**^M^, respectively.

### 3.4 Pathway feature selection and outcome prediction

The iDRW computes the pathway profile based on the multi-layered genegene graph from multiple genomic profiles. The pathway profile is used as an input to the prediction model. We experimented with our proposed method on two types of outcome prediction models: Lasso-Cox regression model to predict OS survival time and the RFE-RFC model to predict metastasis (binary outcome). The prediction performances of pathway profiles obtained from other pathway activity inference methods were evaluated as described for iDRW to achieve a fair comparison.

#### 3.4.1 Lasso-Cox regression model

The Cox proportional hazard model estimates the hazard of each pathway feature at a specific survival time, considering the event status. The regression coefficients represent the degree of correlation of pathway features to the corresponding hazard. We fit a generalized linear model by a maximum likelihood estimation with the li penalty (Lasso), implemented in the R glmnet package (Simon *et al*., 2011). We performed 5-fold crossvalidation in the training set to find the optimal parameter s by choosing the minimum over a grid of λ = 10^Π^, where Π is the sequence decreasing by 0.1 from 10 to −2. The pathway features with non-zero coefficients were selected. We then estimated the hazard of pathways across samples in the test set using the risk scores of the selected pathway features, obtained from training the Lasso-Cox model. In our experiments, the Cox regression model was trained to predict OS adjusted by age, gender, and TNM stage as covariates.

#### 3.4.2 RFE-RFC model

A recursive feature elimination (RFE) is a backward selection algorithm based on the predictors’ importance ranking, and it is a classical and effective method for gene selection (Guyon *et al*., 2002). The algorithm sequentially eliminates less important features based on their ranks. For each iteration, it fits the random forest classification (RFC) model to predict binary outcomes and assesses the importance ranking for predictors. We denote the model as RFE-RFC. As in the Lasso-Cox model, we performed 5-fold cross-validation as a resampling method for important feature selection to reduce the overfitting issue. We fitted the random forest model in the training set, selected the optimal set of pathway features and evaluated the model with the area under the precision-recall curve (AUPRC) in the validation set. The details of the model evaluation are described in the ‘Performance evaluation’ section below. The optimal number of features that should be assessed is also found by the experiments with a varying number of features *N* = [1, 2,⋯, 9,10,15,⋯, 95, 100]. We refitted the random forest model with the optimal features and assessed the prediction performance with AUPRC on the test set. In our experiments, we trained the RFE-RFC model to predict regional lymph node or distant metastasis, adjusted by age and gender as covariates.

### 3.5 Performance evaluation

We randomly split the samples into 70% training and 30% test sets and repeated the process 100 times. To validate each prediction model, we performed 5-fold cross-validation on the training set, which trains the model using four folds (56%) and validates with the remaining one-fold (14%). In the training phase, we fit the optimal model by tuning the hyperparameters by cross-validation and selected the set of pathway features. The fitted model was then evaluated in the test set. As a result, we assessed the performances and the set of selected features after 100 iterations of the entire process for each model in each dataset. The pathways were ranked by their frequencies of being selected for each iteration. Finally, the top-k pathways with more than half frequencies were prioritized.

The Cox regression model that predicts OS was evaluated with a concordance index (C-index). The C-index measures the probability that the observation who is predicted to have a higher risk, has a shorter time-to-event than the other, for a random pair of samples (Harrell *et al*., 1996). The performance of metastasis prediction models was measured with precision-recall (PR) curves. There are three types of binary classification problems: regional lymph node metastasis, distant metastasis, and any metastasis prediction. Precision is the ratio of correctly predicted positive observations of the total predicted positive observations. Recall (Sensitivity) is the ratio of correctly predicted positive observations of all observations in an actual class. PR curves represent the plot of the precision and recall for different thresholds, and the prediction performance is measured with the integral area under the PR curves (AUPRC). Note that we denote the area under the PR curves as AUPRC rather than AUC, as AUC usually refers to the area under the ROC curves. When there is a skew in the class distribution, PR curves provide more accurate performance and better measurements than receiver operating characteristic (ROC) curves (Davis and Goadrich, 2006; Saito and Rehmsmeier, 2015). To address the class imbalance problem, we assigned class weights to the prediction model based on the class distribution, which provides a larger weight on the minority class such that the classifier learns equally from the classes. If the size of the majority class is *N_m_*, and minority class is *N_n_*, we assign the weight of 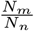 to the minority class. The weight value 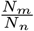 for each prediction model is 3.7 (regional metastasis in BLCA), 5.06 (any metastasis in BLCA), 7.6 (distant metastasis in KIRC), and 8 (any metastasis in KIRC). The classification performances were measured with PR curves. Finally, the median of 100 C-indices or AUPRCs was used as a final performance.

## 4 Results

### 4.1 iDRW contributes to an improved outcome prediction performance

When constructing the multi-layered graph on multiple genomic profiles, we considered four different scenarios (Details are described in ‘Experiments’). We compared the performances of iDRW (ECM) between four scenarios for each prediction model in two cancers, and the difference between the maximum and minimum performance was less than 0.01 (C-index and AUPRC) in any prediction model for both cancers (**Supplementary Table S1**). This result showed that the overall performance was not sensitive to the underlying graph structure. Thus, the iDRW experiments on the multi-layered graph was derived from the scenario (1-a). We evaluated the predictive power of iDRW-based pathway activities for BLCA and KIRC, respectively. We compared the four types of other pathway activity inference models (CORG, PLAGE, DART, and DRW) in each single genomic profile with iDRW for all possible combinations of multiple genomic profiles. Then, we predicted four types of outcomes using the inferred pathway profile as an input: OS, regional lymph node metastasis for distant metastasis-free samples, any metastasis for BLCA; OS, distant metastasis for regional metastasis-free samples, and any metastasis for KIRC.

The prediction results in **Figure 2** showed that pathway activities inferred by DRW mostly performed better than other methods on a single genomic profile when predicting OS in both cancer patient samples. As DART and DRW-based approaches incorporated the significance test results of genes and interaction effects on the graph, they performed better than CORG and PLAGE in all prediction models. The performance improvement of DRW over DART demonstrated that DRW better inferred pathway activities, effectively reflecting the interactions on the graph. When we compared the performances of the DRW-based pathway profile on each genomic profile, we found that the gene expression profile contributed the best to the prediction performances, especially in BLCA, and the methylation profile showed comparable performances in KIRC. We observed that the overall prediction performances on the CNV profile were lower than the other profiles, especially in KIRC, but improvement was observed when we combined it with others using iDRW.

**Fig. 2.**
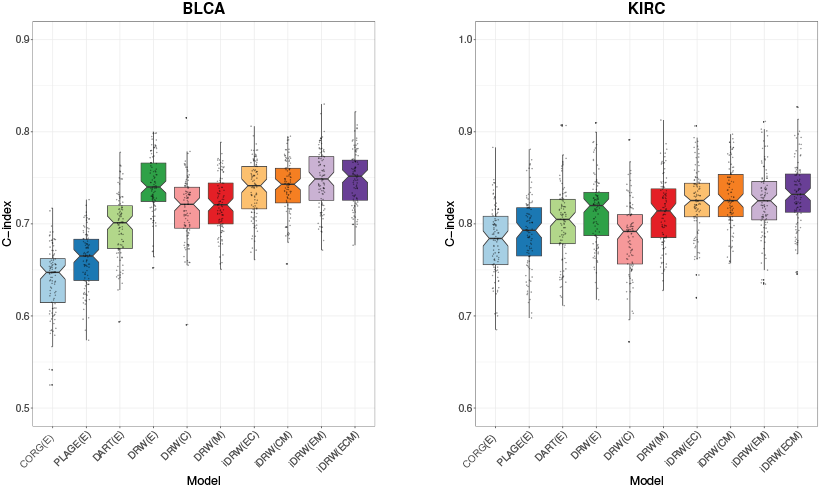
Overall survival (OS) prediction performance comparison between four types of pathway activity inference methods on single genomic profile and iDRW on multiple genomic profiles in bladder cancer (BLCA) and kidney clear cell carcinoma (KIRC). The performance was measured with the median C-index after 100 iterations of the entire process of training and validating models.

Likewise, the metastasis prediction results showed the effectiveness of DRW-based approaches including iDRW on multiple genomic profiles in both cancer datasets. In general, the tendency of performance differences between iDRW and other approaches was similar to the survival prediction results. However, we observed that the contribution to the methylation profile was relatively prominent in kidney cancer metastasis prediction. These results were noticeable in regional lymph node metastasis prediction in BLCA, and in distant or any metastasis prediction in KIRC. The baseline in PR curves of metastasis prediction models corresponded to the proportion of the majority class to the total number of samples, i.e. the junk classifier which predicted with all negatives, denoted as a gray horizontal line in **Figure 3**. In general, the different combinations of integration on three genomic profiles did not significantly differentiate the performances in either cancer. The performances of iDRW on multiple genomic profiles were improved compared to the single genomic profilebased approaches, although it was marginal. These results show that iDRW effectively integrates complementary information, utilizing the interaction effects on the multi-layered network. The prediction performances for each prediction model in both cancer datasets are summarized in **Table 2**.

**Fig. 3.**
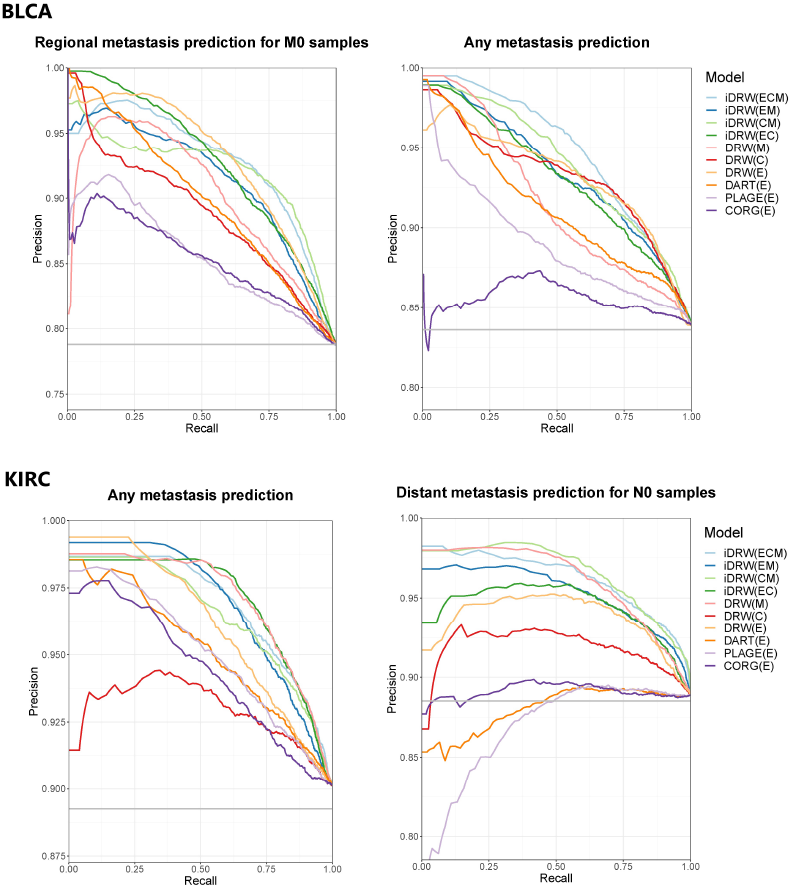
Metastasis prediction performance comparison between four types of pathway activity inference methods on a single genomic profile and iDRW on multiple genomic profiles in bladder cancer (BLCA) and kidney clear cell carcinoma (KIRC). Performance was measured with the area under the precision-recall curves (AUPRC) after 100 iterations of the entire process of training and validating the model.

**Table 2.** Performance summary for each prediction model in bladder cancer (BLCA) and kidney clear cell carcinoma (KIRC)

|  | BLCA |  |  | KIRC |  |  |
| --- | --- | --- | --- | --- | --- | --- |
|  | OS | Regional metastasis | Any metastasis | OS | Any metastasis | Distant metastasis |
|  | C-index* | AUPRC | AUPRC | C-index* | AUPRC | AUPRC |
| CORG(E) | 0.6473 | 0.8531 | 0.8553 | 0.7841 | 0.9445 | 0.8911 |
| PLAGE(E) | 0.6651 | 0.8555 | 0.8903 | 0.7930 | 0.9509 | 0.8684 |
| DART(E) | 0.7012 | 0.8993 | 0.9122 | 0.8048 | 0.9515 | 0.8802 |
| DRW(E) | 0.7400 | **0.9291** | 0.9315 | 0.8199 | 0.9615 | 0.9372 |
| DRW(C) | 0.7213 | 0.8887 | 0.9315 | 0.7918 | 0.9292 | 0.9180 |
| DRW(M) | 0.7208 | 0.9007 | 0.9167 | 0.8141 | 0.9707 | 0.9610 |
| iDRW(CM) | 0.7430 | 0.9207 | 0.9401 | 0.8253 | 0.9650 | **0.9658** |
| iDRW(EC) | 0.7414 | 0.9278 | 0.9309 | 0.8254 | 0.9703 | 0.9442 |
| iDRW(EM) | 0.7488 | 0.9140 | 0.9344 | 0.8251 | 0.9689 | 0.9531 |
| iDRW(ECM) | **0.7564** | 0.9285 | **0.9422** | **0.8294** | **0.9712** | 0.9556 |
| Baseline† |  | 0.7879 | 0.8360 |  | 0.8926 | 0.8852 |

Note that the best prediction performance is emphasized in bold. \*C-index: the median C-index after 100 iterations of the entire process of training and validating the model. †Baseline: the performance when predicting with all negatives, which corresponds to the proportion of the majority class to the total number of samples.

### 4.2 iDRW jointly prioritizes potential driver pathways and genes on multi-omics data

The pathway activities inferred by iDRW were evaluated with prediction models in BLCA and KIRC. Important pathway features were selected for each iteration of training and validating the model. The selected pathways with a frequency of being selected greater than 50 among 100 iterations were prioritized. The complete lists of prioritized features by each prediction model are provided in **Supplementary Table S2** (BLCA) and **S3** (KIRC). The total number of pathway member genes and significantly associated genes with the outcome from the univariate statistical analysis are shown.

**Supplementary Table S4** shows the list of iDRW-prioritized pathways for each prediction model in both cancer datasets. To demonstrate the effectiveness of iDRW when prioritizing pathways, we showed pathways only when we integrated multiple genomic profiles on the graph, not in single omics-based approaches. The best-performing combinations of iDRW models are shown. The results showed that iDRW identified 15 significantly associated pathways with bladder cancer prognosis. The pathways are categorized into the main- and sub-class derived from the KEGG pathway database, as shown in **Supplementary Table S4**. Forty-two pathways were found to be related to regional or any metastasis in BLCA. Most of the dominant pathways were related to metabolism or organismal systems such as the digestive system. Interestingly, 11 human disease pathways were identified in bladder cancer metastasis, and 7 out of 11 pathways were related to infectious diseases, such as bacterial, parasitic, and viral diseases. Additionally, 4 immune system-related pathways and others including neurodegenerative disease and microRNAs in cancer pathways were found in bladder cancer metastasis. There were a relatively small number of pathways in KIRC compared with BLCA. iDRW identified 6 pathways related to kidney cancer prognosis. There were 12 distant metastasis-associated pathways in KIRC, and 9 out of 12 pathways were related to metabolism. The one bacterial infectious disease pathway (pertussis) was identified in any metastasis for KIRC.

In addition to the iDRW-prioritized pathways, we obtained the pathways that were important both in single- and multi-omics profiles, which appeared more than three (count>3) among seven DRW-based models: DRW on **G**^E^, **G**^C^, and **G**^M^; iDRW on **G**^EC^, **G**^EM^, **G**^CM^, and **G**^ECM^, as shown in **Supplementary Table S5** (The frequently appeared pathways were emphasized as bold). There were 10 and 4 commonly identified pathways to both single- and multi-omics based models in BLCA and KIRC, respectively. They include 5 pathways related to bladder cancer prognosis. Five pathways were found to be associated with bladder cancer metastasis, mostly regional metastasis. We found 4 metabolism pathways related to kidney cancer prognosis or distant metastasis, such as glycan biosynthesis and metabolism, and lipid metabolism.

In summary, the most dominant pathways were related to metabolism both in iDRW-prioritized and commonly identified pathways. The metabolism, infectious disease, and digestive or endocrine system-related pathways were specifically found using iDRW. We observed that the commonly important pathways were mostly associated with metabolism, cell growth and death, and human diseases (not infectious disease). Especially in human disease pathways, infectious disease pathways were found only in iDRW-prioritized pathways, and there were mostly related to regional metastasis in BLCA, e.g., toxoplasmosis pathway (KEGG: map05145), malaria (KEGG: map05144), leishmaniasis (KEGG: map05140). We found that both the iDRW-prioritized and commonly identified pathways were associated with cancer survival or metastasis. The evidences were shown in **Supplementary Discussion**.

### 4.3 iDRW facilitates integrative gene-gene network analysis

We visualized the top-10 pathways prioritized by iDRW on the multilayered network **G**^ECM^ for the OS prediction model in BLCA. The genes that were significantly associated with bladder cancer prognosis within pathways from three genomic profiles were jointly analyzed on the network. The graph was formally constructed by assigning between-layer edges from all pairwise combinations of genes in different layers, but 20-25 edges that were randomly chosen between each pair of different layers are shown for visualization. The size of nodes represents the log-transformed *p*-value of significant genes, i.e., the larger the node size, the higher is the significance level of the gene. We differentiated each node into different colors and shapes according to the type of genomic profile.

As shown in **Figure 4**, there was an exclusively large number of significant genes related to alcoholism (*N* =67), necroptosis (*N* =54), and neuroactive ligand-receptor interaction pathways (*N* =77); the last one was ranked 12^th^ by frequency. The results showed that these three pathways and genes greatly contributed to bladder cancer prognosis. The 36 histone genes, which were mostly included in the largest cluster HIST1, were found in the alcoholism or necroptosis pathway, the biggest pathways among the top-10 pathways. Overall, the most significant genes on average were from the methylation profile; ARSB (methylation), ALOX15 (methylation), CPT1B (gene expression), ITGB7 (gene expression), ABCA (methylation), and MAPK3 (methylation). The phototransduction, necroptosis, and intestinal immune network for IgA production pathways were prioritized specifically by iDRW. There were 5 commonly identified pathways both in single- and multi-omics profiles: ubiquitin-mediated proteolysis pathway, fatty acid metabolism pathway, ABC transporters pathway, glycosaminoglycan degradation pathway, and phenylalanine, tyrosine, and tryptophan biosynthesis pathway.

**Fig. 4.**
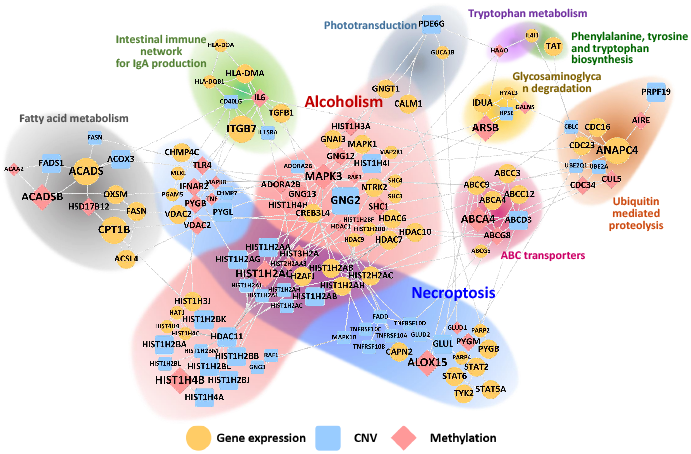
Integrated pathway-based gene-gene network using the top-10 pathways prioritized by iDRW (ECM) for OS prediction in BLCA. We note that the node size represents the log-transformed *p*-value of significant genes, and the node color and shape represent the type of genomic profiles; gene expression profile (yellow ellipse), CNV profile (blue round rectangle), methylation profile (red diamond).

## 5 Discussion

In this study, we proposed the multi-layered network-based pathway activity inference method on multi-omics data (iDRW) and experimentally validated the method with two types of clinical outcome prediction models for urologic cancer integrative analysis. The main advantages of our framework are summarized as follows. First, it is generally applicable to any type of data that integrates multi-omics data on the pathwaybased network while other competitive methods targeted a single genomic profile. Second, as it is not sensitive to the underlying graph structure, the researchers can customize multi-layered network. Finally, it facilitates the integrative network-based pathway-level analysis: pathway activity pattern analysis, outcome-related pathways prioritization, and integrated network visualization. The experimental results showed that the method not only contributes to an improved outcome prediction performance but also provides better biological insights into the pathways and genes prioritized by the model from a comprehensive perspective.

The marginal performance improvements in iDRW may have been affected by the incomplete mapping of genes within pathways. In our experiments, approximately 30% of genes in genomic profiles were mapped on average in the pathway-based gene-gene graph using the KEGG pathway database. This issue can be resolved through complete mapping by merging with another existing pathway database, such as Reactome, BioPAX, and WikiPathways. In addition, the performance improvement of iDRW in metastasis prediction was insignificant, even though we alleviated the class imbalance problem in the metastasis prediction model by giving class weights when training the model. As iDRW contributed to an improved OS prediction power, we expect a more improved classification performance, given the balanced class distribution. We identified that iDRW showed the complementary effect when integrating multi-omics data with large variances, as shown in **Supplementary Figure S2**. These results demonstrated that iDRW robustly combines heterogeneous data, reducing the noise.

Furthermore, we visualized and analyzed the multi-layered gene-gene network for the integrative urologic cancer analysis. Since cancer is a complex disease caused by genetic and/or epigenetic changes at different molecular levels such as the DNA sequence, expression, methylation, copy number variation, metabolite, and proteome, the effective multi-omics data integration framework is essential to understand the complex nature of cancer biology. Based on the prioritized pathways, genes and genegene network provided by our framework, we might be able to provide a holistic understanding of the pathophysiological mechanisms in cancer development and progression. It is possible to identify novel biomarkers for cancer diagnosis and prognosis, provide risk prediction of cancer patients, and discover efficient targeted anti-cancer agents. Future works are still needed that include pan-cancer analysis and integration with other omics data such as proteomics and metabolomics.

## Acknowledgements

We gratefully acknowledge the TCGA Consortium and all its members for the TCGA Project initiative, for providing sample, tissues, data processing and making data and results available. The results published here are in whole or part based upon data generated by The Cancer Genome Atlas pilot project established by the NCI and NHGRI. Information about TCGA and the investigators and institutions that constitute the TCGA research network can be found at http://cancergenome.nih.gov.

