## Supplementary Materials for "Multi-layered network-based pathway activity inference using directed random walks: application to predicting clinical outcomes in urologic cancer"

### 1 Supplementary Materials

#### 1.1 Supplementary Discussion

In our experiments, the gene expression profile contributed the best to bladder cancer survival or metastasis prediction as in previous studies [Maeda et al., 2018, Seiler et al., 2016]. The results showed that the methylation profile plays an important role in kidney cancer prognosis or metastatic progression. Several studies suggest that DNA methylation changes are one of the promising biomarkers for prognosis and progression of kidney clear cell carcinoma [Lasseigne and Brooks, 2018]. For example, it has been discovered that a DNA methylation biomarker can differentiate kidney tumor from benign adjacent kidney tissue, irrespective of the tumor histology [Lasseigne et al., 2014]. In addition to the gene expression profile, the performances of CORG, PLAGE, DART, and DRW on CNV or the methylation profile are shown in **Supplementary Figure S1**. In general, performance improvement was observed in DRW. However, the performance difference between DRW and DART was minimal on the methylation profile. As shown in **Supplementary Figure S2**, DRW-inferred pathway activities close to the minimum were predominant on the methylation profile in comparison to that inferred by DART. The DRW infers pathway activities by combining the normalized expression values, statistical significance score, and weight of genes within pathways by the product of the random walk procedure. As DART utilizes the expression values and statistical significance of genes, the random walk on the graph with methylation features might have degraded the performance, by the negative effect of considering the interactions between methylation features. This phenomenon has been observed only in bladder cancer survival prediction models. The impact of DNA methylation features on kidney cancer prognosis might have compensated for this performance degradation.

iDRW infers pathway activities across whole samples and the pathway profile is used as an input to the prediction model. We analyzed the iDRW-inferred pathway activity patterns across two urologic cancers for each predictive outcome. The heatmap shows the pathway activity pattern with genes that were significantly associated with OS or metastasis (**Supplementary Figure S3**). Note that regional lymph node metastasis was used in BLCA and distant metastasis in KIRC. The intersection of top-$k$ prioritized pathways between BLCA and KIRC is shown; $k = 80$, which is experimentally set for better visualization. Each pathway profile was normalized to have a range between zero and one. The pathways and samples were clustered via Ward's hierarchical agglomerative clustering method [Murtagh and Legendre, 2014].

In total, there were 20 OS-related and 26 metastasis-related pathways, which are commonly identified in both cancers. The distinctive pathway activity patterns were observed across cancer types both in OS and metastasis. Two large clusters of pathways related to OS were observed, which could be clearly distinguished by cancers. The heatmap shows that the pathway activities in the largest cluster were mostly low, close to the minimum in BLCA and relatively high in KIRC, and the opposite was observed in the second largest one. The largest one contained 12 pathways, including 5 genetic information processing pathways, 4 metabolism pathways, 2 human disease pathways (immune disease and substance dependence), and one sensory system pathway. The second largest one contained 8 pathways: four metabolism-related pathways, a digestive system pathway, an excretory system pathway, an endocrine and metabolic disease pathway, and a signaling molecules and interaction pathway. We observed that genetic information processing-related pathways such as replication and repair, translation, and transcription were only included in the largest cluster. Most of the pathways in **Supplementary Figure S3** showed cancer-specific characteristic, which is quite distinguishable.

Likewise, two large clusters were present in relation to regional or distant metastasis. The largest cluster showed an extremely distinctive pattern, the activities of which are relatively high, close to the maximum in distant metastasis (KIRC) and low in regional metastasis (BLCA). They included 6 metabolism pathways, 3 organismal systems pathways, 3 human disease pathways (2 of 3 related to infectious diseases), 3 signal transduction pathways, and one cellular process pathway. In contrast, the second largest one shared characteristics between cancers. It included 4 metabolism pathways (3 of 4 for cofactors and vitamins), and 6 other pathways (e.g., immune system, environmental adaption, endocrine, and metabolic disease). Following the results of iDRW-prioritized pathways related to regional metastasis (**Supplementary Table S4**), infectious disease pathways were found only in the largest cluster, which is related to metastasis, e.g., salmonella infection pathway (KEGG: map05132) and influenza A pathway (KEGG: map05164). This finding implies that the infectious disease-related genes play a role in urologic cancer metastasis as a risk factor.

In total, 22 metabolism-related pathways out of 69 iDRW-prioritized pathways were found to be associated with both bladder and kidney cancer survival or metastasis. Metabolism pathways, including amino acid, lipid, nucleotide, carbohydrate, and glycan biosynthesis and metabolism play crucial roles in the prognosis and metastasis of urologic cancers [Chen et al., 2017, Ohyama, 2008, Weiss, 2018]. We found several lines of evidence concerning associations between three environmental information processing pathways and bladder cancer prognosis or progression. Cell adhesion molecules and their related pathways are known to be strong candidate targets for the inhibition of cancer progression, including bladder cancer [Bryan, 2015]. Cytokine-cytokine receptor interaction and neuroactive ligand-receptor interaction pathways are signaling pathways that have been shown to be associated with bladder cancer progression by the enrichment analysis in this study [Fang et al., 2013]. Nicotine increases tumor growth and induces acquired chemoresistance through activation of the PI3K/Akt/mTOR pathway in bladder cancer [Yuge et al., 2015]. Following the results of this study, nicotine addiction, nicotinate, and nicotinamide metabolism, as well as PI3K-Akt signaling pathway were shown to be risk factors for bladder cancer prognosis and regional metastasis. MicroRNAs in the cancer pathway were found only in the regional metastasis prediction model in BLCA. The miRNA signatures of cancer were observed in various cancer studies, including bladder cancer [Enokida et al., 2016, Xie et al., 2017]. The dysregulation of miRNAs has been shown to be closely related to the pathological features of bladder cancer, such as the tumor stage, grade, metastasis, recurrence and chemo-sensitivity [Dong et al., 2017].

Eleven immune system-related pathways, including infectious disease and immune disease, were related to regional lymph node metastasis in bladder cancer. The four immune system or disease-related pathways were commonly identified both in bladder and kidney cancer, including complement and coagulation cascades, NOD-like receptor signaling pathway, salmonella infection, and influenza A. It is widely known that chronic inflammation has serious effects in the initiation and progression of cancer metastasis, and the immune system plays a significant role in the metastatic progression of tumors [van der Horst et al., 2012, Arai et al., 1981]. We found several bacterial, parasitic, and viral infectious disease-related pathways related to the risk of bladder cancer regional metastasis, including malaria, tuberculosis, toxoplasmosis, hepatitis C, leishmaniasis, African trypanosomiasis, and pertussis. The studies showed evidence that parasitic infection, including schistosomes, trematodes, and toxoplasmosis causes urinary bladder cancer progression [Dyck and Mills, 2017, Thun et al., 2004]. In addition to those infectious or immune disease pathways, several digestive systems pathways were found to be risk factors for bladder cancer regional metastasis, e.g., bile/pancreatic secretion, cholesterol, lipid and fatty acid metabolism, fat digestion, and absorption. They have been mostly shown to be associated with gastrointestinal cancer, demonstrating no strong associations with regional lymph node metastasis in bladder cancer, which are novel findings in iDRW.

We also found evidence for the iDRW-identified potential driver pathways for KIRC survival or distant metastatic progression. The effect of the dysregulation of metabolic pathways, including fatty acid, lipid, carbohydrate, amino acid, glycan metabolism pathways, on renal cell carcinoma has been studied and the key metabolic abnormalities underlying renal cancer carcinogenesis has been analyzed [Massari et al., 2015, Rathmell et al., 2018]. In accordance with our findings, tryptophan metabolism has been shown to be an important pathway in renal cancer [Liu et al., 2019]. Hepatocyte nuclear factor-1beta (HNF1$\beta$) gene mutations cause maturity-onset diabetes of the young type 5 (MODY5), and the expression of HNF1$\beta$ has been shown to be associated with cancer risk in several tumors, including renal cancer [Yu et al., 2015, Verhave et al., 2016]. Renal damage due to cisplatin toxicity is prevented to a great extent by the anti-inflammatory effect of oxytocin [Erbas et al., 2014], which shows the possible association between the oxytocin signaling pathway and renal cancer prognosis. There is a possible connection between the taste transduction pathway and renal clear cell carcinoma, wherein several factors, including renal dysfunction, may affect taste perception, which is related to the advanced cancer state [Murtaza et al., 2017]. The effect of morphine treatment on cancer pain have been widely studied, and the side effects and toxicity of opioid use have been studied, leading to the development of renal failure, and kidney disease [Mallappallil et al., 2017]. Pathways related to vitamin supply were identified, including vitamin digestion and absorption and biotin metabolism. Several *in-vitro* and *in-vivo* studies have shown that vitamin D inhibits renal cell cancer cell proliferation, angiogenesis, clonogenicity, and metastasis, induces cell differentiation and prolongs survival [Blomberg Jensen et al., 2010, Fujioka et al., 2000, Nagakura et al., 1986]. Mutations in some FANC genes, among which FAN1 is included as a significant gene in the Fanconi anemia pathway, cause Fanconi anemia and significantly increase cancer susceptibility sporadically in the general population [Niraj et al., 2019]. FAN1 mutations have been shown to cause chronic kidney disease [Zhou et al., 2012].

We note that cancer patients with distant metastasis, but without regional lymph node metastasis (any T/N0/M1), are quite rare in most cases. There is only one case in BLCA, but 28 cases in KIRC in particular. The experiments in our study revealed prognostic risk factors for those exclusive cases. We found evidence for the association between the risk pathway candidates and the risk of renal cancer, e.g., vitamin-related pathways, Fanconi anemia, caffeine metabolism, and several metabolism pathways. Studies to reveal the clinical association between them and distant metastasis should be conducted to underpin those novel findings.

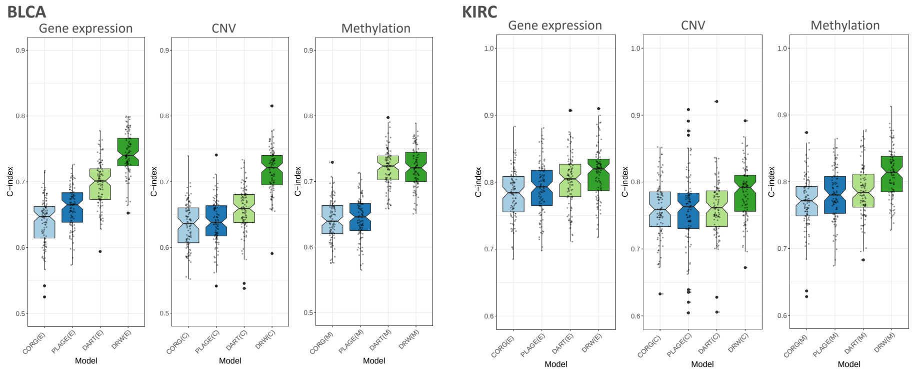


Figure 1: Overall survival (OS) prediction performance comparison between four types of pathway activity inference methods for each genomic profile; gene expression, CNV, methylation profile in bladder cancer (BLCA) and kidney clear cell carcinoma (KIRC). Performance was measured with the mean C-index after 100 iterations of the entire process of training and validating the model

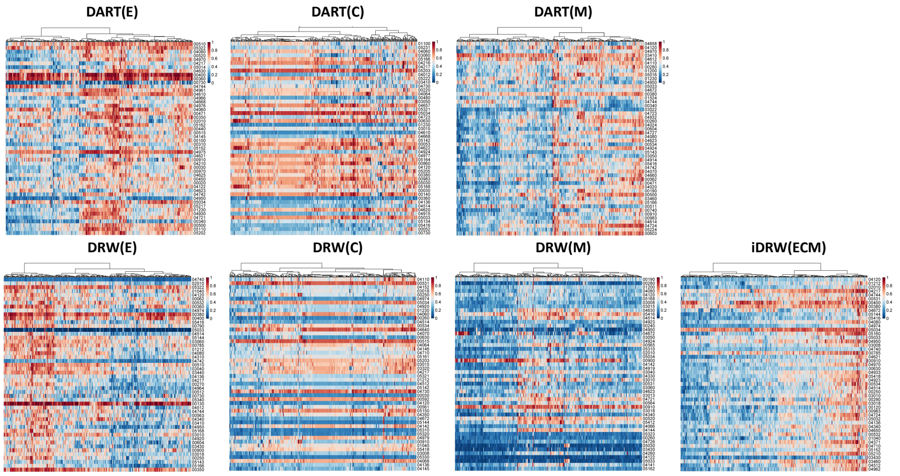


Figure 2: Heatmap of pathway activities inferred by DART and DRW for each genomic profile, and iDRW (ECM). The significance of genes associated with OS are measured with univariate cox regression analysis for each pathway. The pathway profile was normalized to have a range between zero and one. The rows and columns represent pathways and samples, respectively, and columns (samples) are clustered via Ward's hierarchical agglomerative clustering method. Note that rows (pathways) are not clustered to compare the pathway activity patterns.

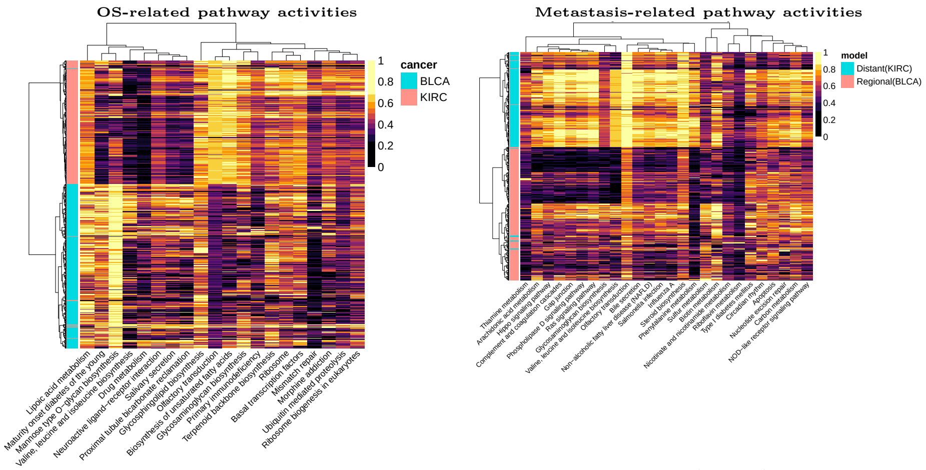


Figure 3: Heatmap of pathway activities inferred by iDRW(ECM). The significance of genes associated with OS (left) or metastasis (right) was measured with univariate cox regression analysis or a two-tailed t-test for each pathway. The pathway profile was normalized to have a range between zero and one. The rows and columns represent samples and pathways respectively, which are clustered via Ward's hierarchical agglomerative clustering method.

### 2 Supplementary Tables

Supplementary tables are provided in excel file.
